## Supplementary Figures for "An integrated proteome and transcriptome of B cell maturation defines poised activation states of transitional and mature B cells"

### Supplementary data

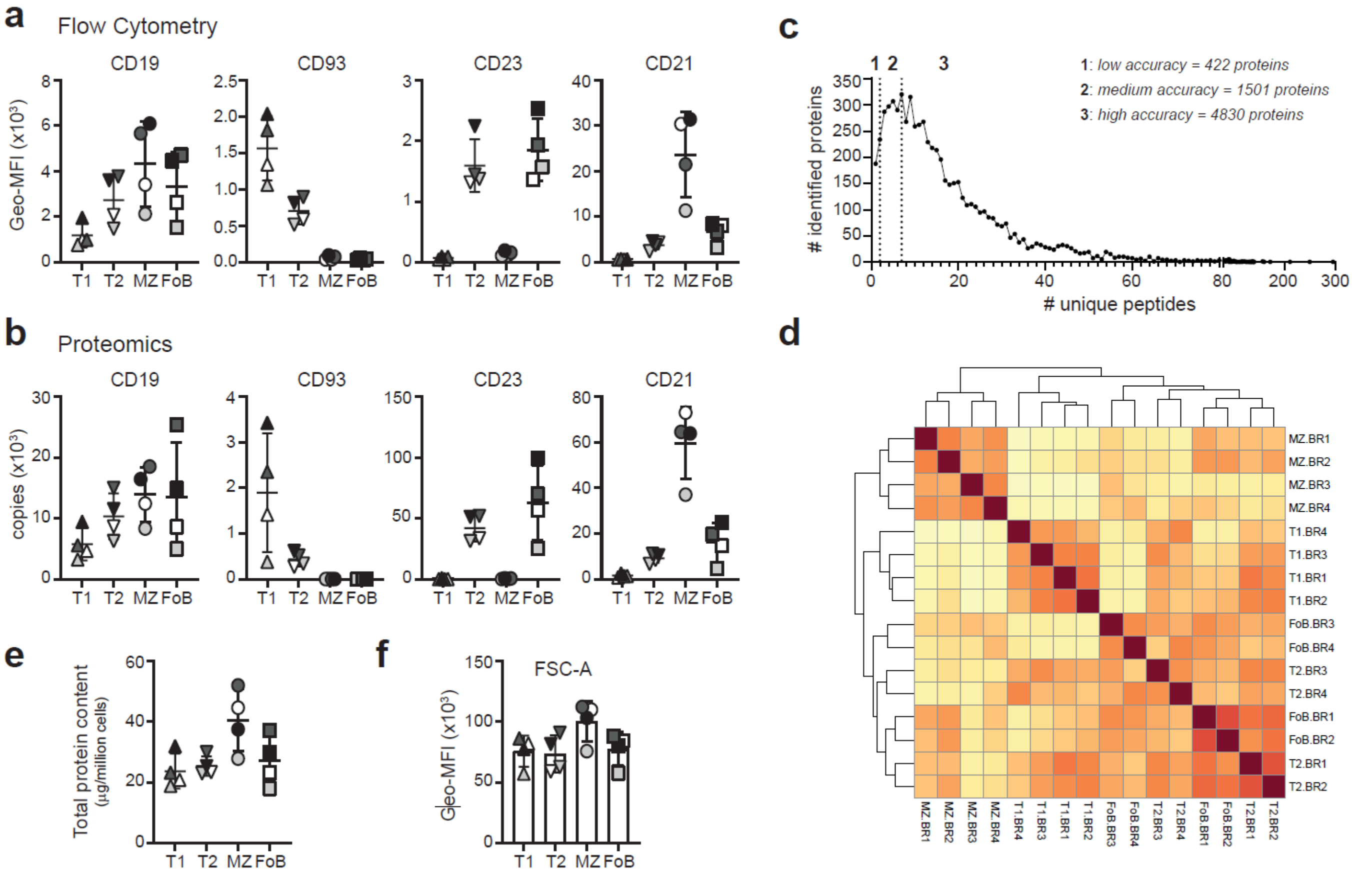


**Supplementary Figure 1: QC of proteomic analysis. a-b**, CD19, CD93, CD23/FcER2 and CD21/CR2 protein content of T1, T2, MZ and FoB cells as measured by flow cytometry and displayed as geometric fluorescence intensity (Geo-MFI) (**a**) or as measured by mass spectrometry and calculated using the proteomic ruler approach (**b**). **c**, Distribution of identified proteins in relation to the number of their assigned unique peptides. Graph displays only 6,753 protein groups were found in at least three of four biological replicates. Full dataset is reported in Table S1. **d**, Heatmap visualizing the Pearson correlation coefficients between the log10-transformed copy numbers for each B cell population and biological replicate. **e**, Total protein content of T1, T2, MZ and FoB cells calculated by proteomics. **f**, Geometric-mean fluorescence intensity (Geo-MFI) of forward light scatter area (FSC-A) of B cell subsets as detected during flow cytometric sorting. In **a**, **b**, **e**, **f** each biologically independent sample was indicated using a greyscale (n= 4 mice; mean ±SD).

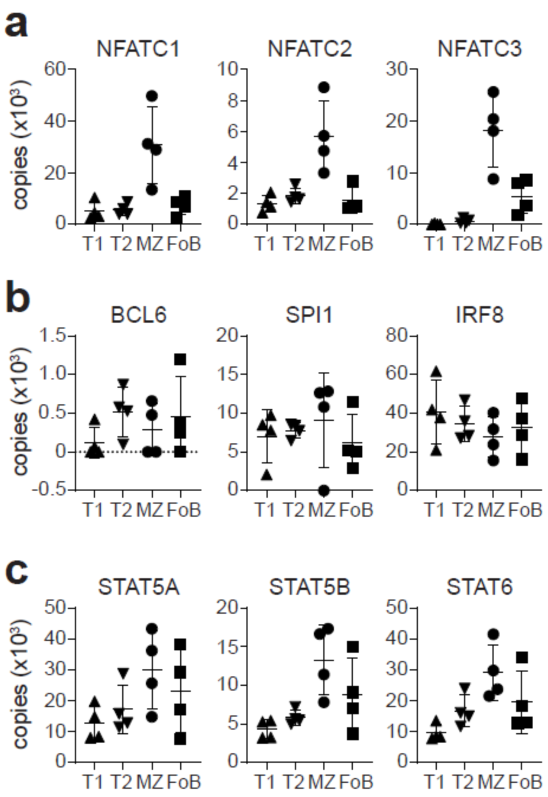


**Supplementary Figure 2: Quantitation of selected transcription factors in B cell subsets. a-c**, Copy numbers of activation-related NFATC-1, -2, -3 (**a**), germinal-centre related BCL6, SPI1, IRF8 (**b**), and cytokine-related STAT5a, STAT5b, STAT6 (**c**) transcription factors in T1, T2, MZ and FoB cells (n= 4 mice; mean ±SD).

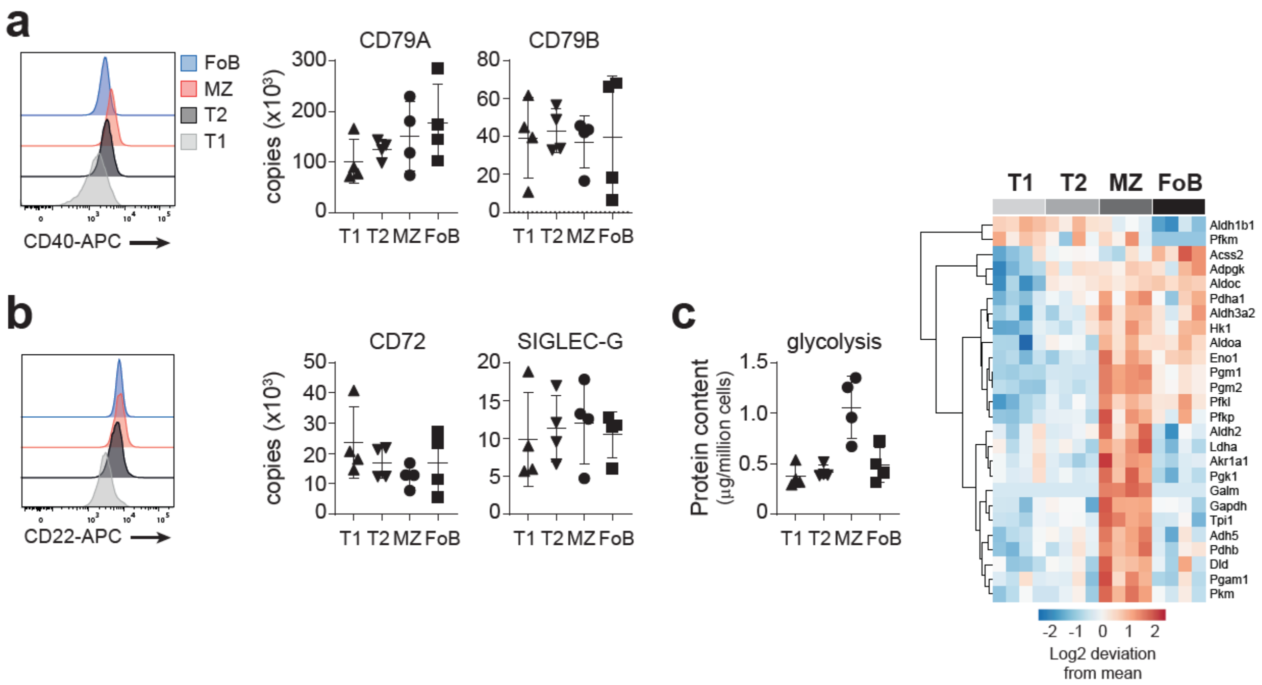


**Supplementary Figure 3: BCR- and glycolysis-associated proteins in B cell subsets. a**, **b**, Histograms depict CD40 (**a**) and CD22 (**b**) expression by flow cytometry. Graphs show mean ±SD of protein copy numbers of CD79a and CD79b (**a**), and CD72 and SiglecG (**b**). **c**, Left: protein content of detected components of the glycolytic pathway (Kyoto Encyclopedia of Genes and Genomes map00010). Right: heat map showing the log2-fold deviation from the mean of normalized copy numbers of 38 glycolysis-related proteins that were differentially expressed among T1, T2, MZ and FoB cells.

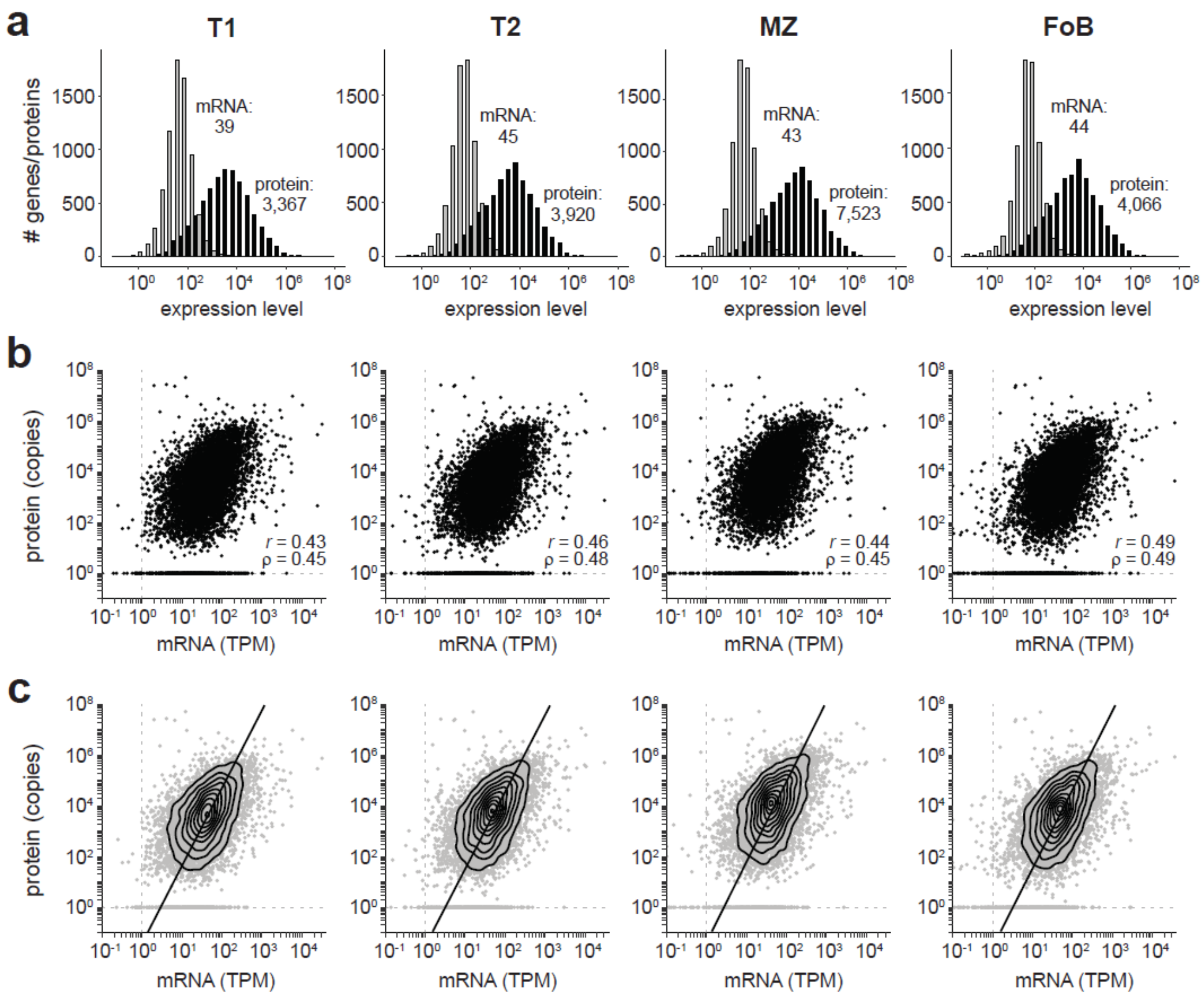


**Supplementary Figure 4: Across-gene comparison of transcriptomic and proteomic data. a**, Histograms indicate the distribution of mRNA (grey) and protein (black) abundance in T1, T2, MZ and FoB cells. Numbers indicate median of TPMs and protein copy numbers, which was calculated on 7,303 genes that were detected by both Illumina sequencing and proteomics. The full list of TPMs and copy numbers is provided in Table S4. **b**, Scatter plots show the average mRNA abundance (TPM) and the average protein abundance (copies) of 7,303 genes that were detected in T1, T2, MZ and FoB cells. Pearson correlation coefficients (*r*) and Spearman correlation coefficients (*ρ*) were calculated for each B cell subset. **c**, Density contours (black) overlaying scatter plots as in b (here reported in grey) were used to calculate a trend line that assumes a positive relationship between mRNA and protein expression in each B cell subset.

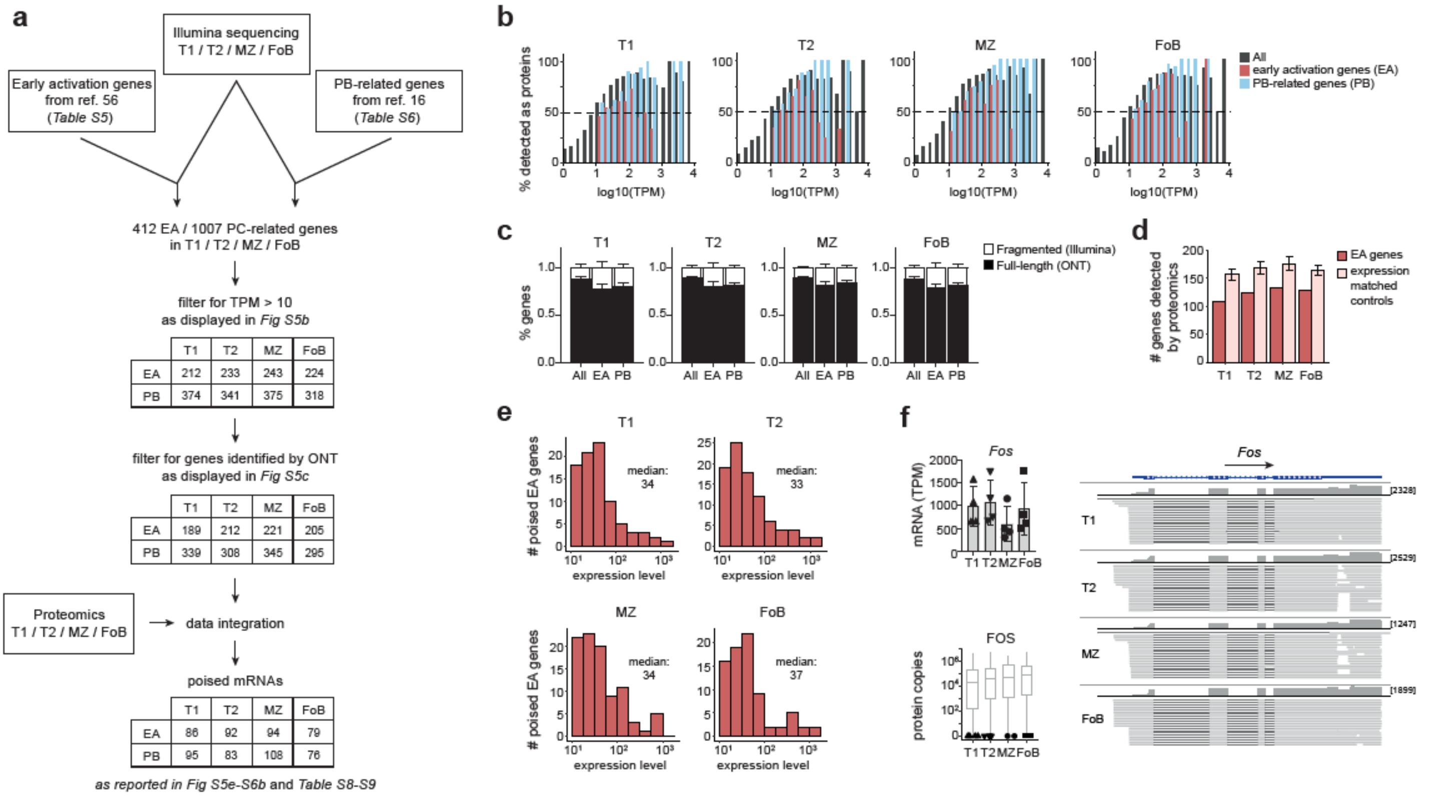


**Supplementary Figure 5: Selection of poised mRNAs in B cells. a**, Analysis workflow for the identification of poised early activation (EA) and PB-related (PB) genes in T1, T2, MZ and FoB cells. **b**, Percentage of detected proteins at indicated TPMs of protein-coding mRNAs was calculated for each B cell subset. Bar graphs show the overall dataset as measured by Illumina sequencing (dark grey), and selected early activation genes (red) and PB-related genes (blue) with TPM > 10. **c**, Graphs indicate the proportion of genes as in **b**) that were detected in full-length (black) by ONT sequencing. **d**, Number of genes detected at the protein level for the 189-221 full-length protein-coding early activation genes found in the transcriptome of T1, T2, MZ and FoB cells, compared with expression-matched controls. For expression-matched controls, the bar represents the median across 100 control gene sets, and the error bars the 5$^{th}$ and 95$^{th}$ percentiles. A difference with $p < 0.1$ is found when the number of proteins detected within early activation genes is lower than the error bars of the corresponding control gene sets. **e**, Histograms indicate the distribution of poised early-activation mRNA abundance in T1, T2, MZ and FoB cells. Numbers indicate median of TPMs as calculated by Illumina sequencing. All genes are listed in Table S8. **f**, Left top: *Fos* TPM as measured by Illumina sequencing. Left bottom: Black symbols show FOS protein detection within our proteomic dataset. Grey boxes represent protein copy numbers of expression-matched controls. Right: Genome browser view displays individual long-reads from ONT sequencing data of mouse *Fos* (chr12: 85,473,813-85,477,428) visualized with the Integrative Genomics Viewer. RefSeq GRCm38 annotation (blue), coverage of (dark grey) and aligned long-reads (light grey) were reported for each gene. Lines connecting light grey boxes indicate splicing junctions between aligned sequences. In squared brackets the maximum read coverage for each B cell population.

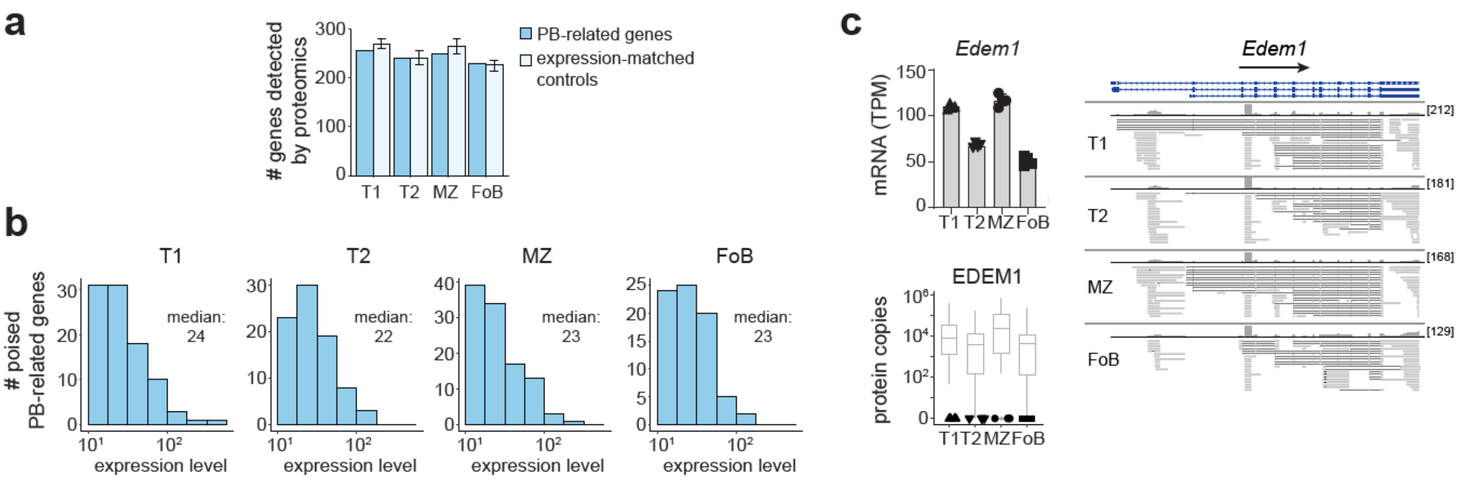


**Supplementary Figure 6: Selection of PB-related genes in B cells. a**, Number of genes detected at the protein level for the 295-345 full-length PB-related genes found in the transcriptome of T1, T2, MZ and FoB cells, compared with expression-matched controls. For expression-matched controls, the bar represents the median across 100 control gene sets, and the error bars the 5$^{th}$ and 95$^{th}$ percentiles. A difference with $p < 0.1$ is found when the number of proteins detected within PB-related genes is lower than the error bars of the corresponding control gene sets. **b**, Histograms indicate the distribution of poised PB-related mRNA abundance in T1, T2, MZ and FoB cells. Numbers indicate median of TPMs as calculated by Illumina sequencing. All genes are listed in Table S9. **c**, Left top: *Edem1* TPM as measured by Illumina sequencing. Left bottom: Black symbols show EDEM1 protein detection within our proteomic dataset. Grey boxes represent protein copy numbers of expression-matched controls. Right: Genome browser view displays individual long-reads from ONT sequencing data of mouse *Edem1* (chr6:108,828,338-108,859,976) visualized with the Integrative Genomics Viewer. RefSeq GRCm38 annotation (blue), coverage of (dark grey) and aligned long-reads (light grey) were reported for each gene. Lines connecting light grey boxes indicate splicing junctions between aligned sequences. In squared brackets the maximum read coverage for each B cell population.

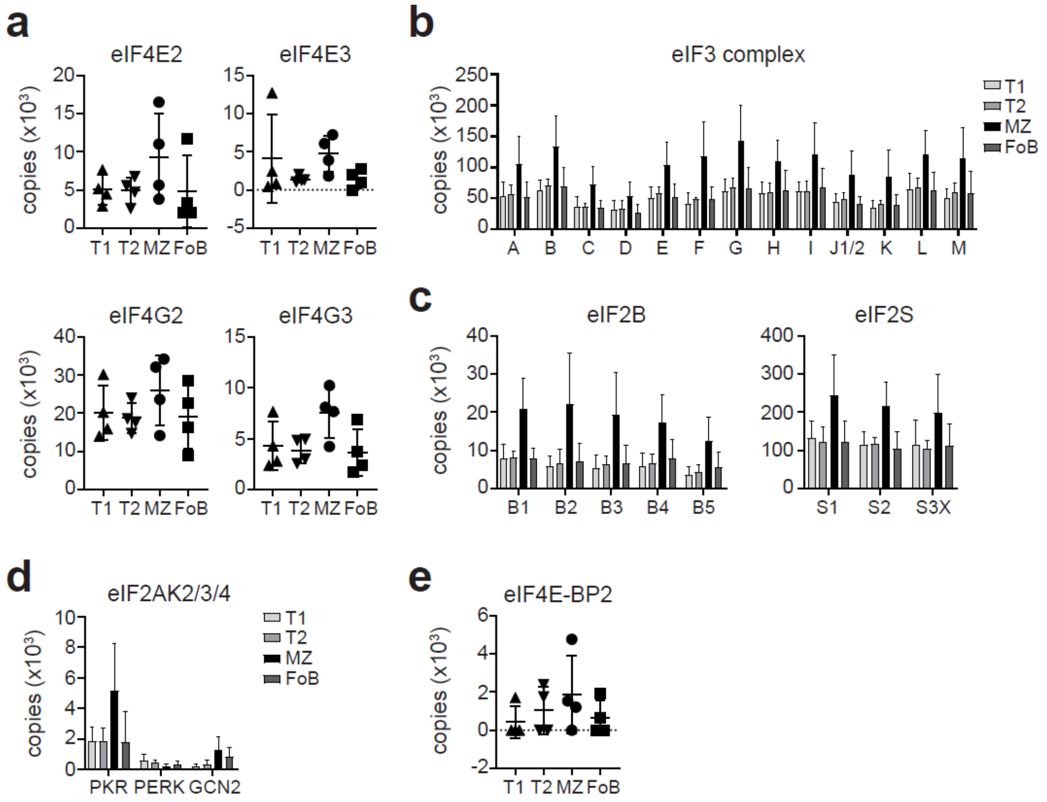


**Supplementary Figure 7: Quantitation of factors promoting or inhibiting translation initiation in B cell subsets. a-d**, Copy numbers of components of the eIF4F translation initiation complex (**a**), the eIF3 complex (**b**), the eIF2 complex (**c**), eIF2α kinases (**d**), and eIF4E-binding protein 2 (**e**).

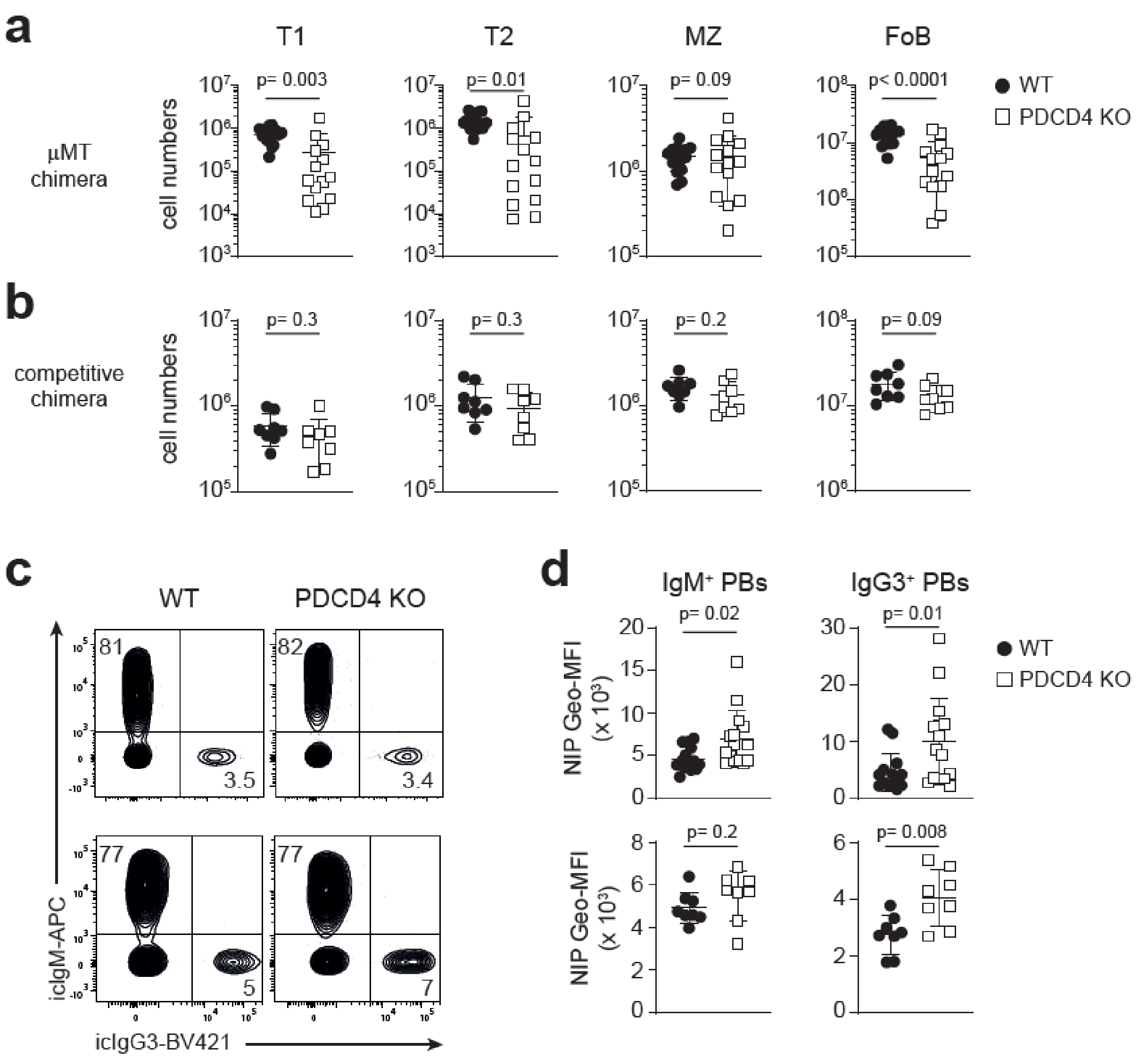


**Supplementary Figure 8: Reconstitution and immunization of WT and PDCD4 KO BM chimeras.** **a**, **b**, Absolute numbers of CD45.2$^+$ T1 (CD19$^+$ CD93$^+$ IgM$^+$ CD23$^-$), T2 (CD19$^+$ CD93$^+$ IgM$^+$ CD23$^+$), MZ (CD19$^+$ CD93$^-$ CD21$^+$ CD1d$^+$ CD23$^-$) and FoB (CD19$^+$ CD93$^-$ CD21$^-$ CD1d$^-$ CD23$^+$) cells from spleen of □MT chimeras (**a**) or competitive chimeras (**b**) 8-9 weeks after BM reconstitution. **c**, Representative contour plots of intracellular-IgM and intracellular-IgG3 expression of CD138$^+$ TACI$^+$ CD19$^{int/low}$ IgD$^-$ CD45.2$^+$ PBs. **d**, Geometric-mean fluorescence intensity (Geo-MFI) of NIP$^+$ intracellular-IgM and intracellular-IgG3 PBs as in **c**. In **c-d** top row shows □MT chimeras, whereas bottom row shows competitive chimeras seven days after NP-Ficoll immunization. Graphs in **d** indicate mean ±SD of n= 14-15 mice for µMT chimeras; n= 8 mice for competitive chimeras. Unpaired Student's *t*-test was performed between WT (black circles) and PDCD4 KO (open squares) samples.

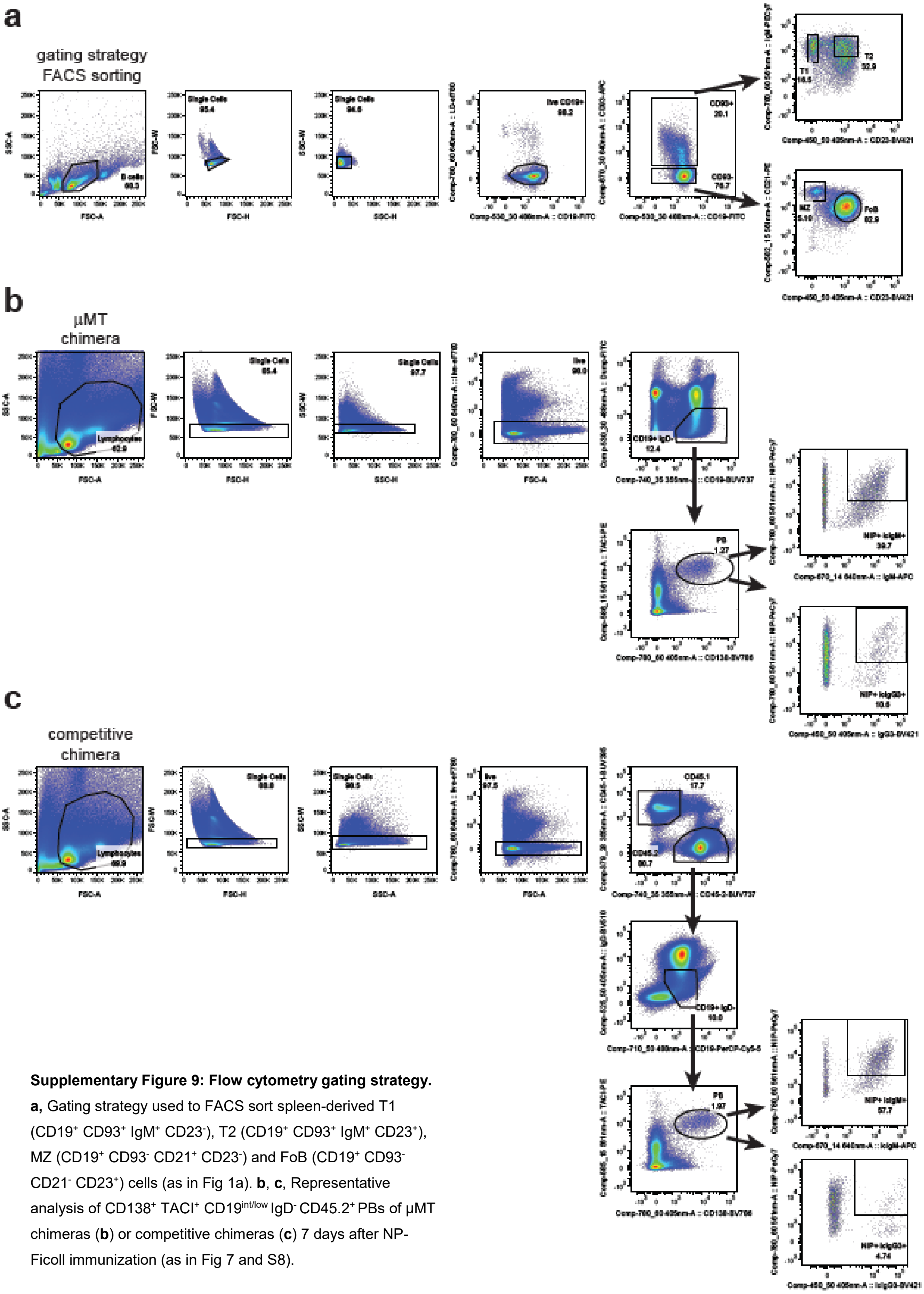


**Supplementary Figure 9: Flow cytometry gating strategy.** **a,** Gating strategy used to FACS sort spleen-derived T1 (CD19$^{+}$ CD93$^{+}$ IgM$^{+}$ CD23$^{-}$), T2 (CD19$^{+}$ CD93$^{+}$ IgM$^{+}$ CD23$^{+}$), MZ (CD19$^{+}$ CD93$^{-}$ CD21$^{+}$ CD23$^{-}$) and FoB (CD19$^{+}$ CD93$^{-}$ CD21$^{-}$ CD23$^{+}$) cells (as in Fig 1a). **b**, **c**, Representative analysis of CD138$^{+}$ TACI$^{+}$ CD19$^{int/low}$ IgD$^{-}$ CD45.2$^{+}$ PBs of µMT chimeras (**b**) or competitive chimeras (**c**) 7 days after NP-Ficoll immunization (as in Fig 7 and S8).
